# Casanovo-TMT: deep learning-based de novo sequencing of TMTpro-labeled peptides and phosphopeptides

**DOI:** 10.64898/2026.11.15.766634

**Authors:** Han Byeol Kim, Kwang-eun Kim

## Abstract

De novo peptide sequencing reads peptides directly from tandem mass spectra, but none of the released deep learning models can sequence peptides labeled with isobaric tags such as TMTpro, which are widely used for multiplexed quantitative proteomics. We extended the residue vocabulary of Casanovo with TMTpro tags and phosphorylation, fine-tuned it on 21 million spectra from our laboratory, most of them TMTpro-labeled, and re-ranked the candidates in the beam by their agreement with the measured precursor mass. The resulting model, Casanovo-TMT, predicted the exact peptidoform for 84.7% of the TMTpro-labeled spectra in our test set and for 87.4 and 79.6% of the spectra in two public global-proteome datasets from other laboratories, whereas the extended vocabulary without fine-tuning gave 0.3%. Discarding the lowest-scoring predictions raised precision to 95% while retaining 81.7% of the test spectra. Casanovo-TMT also predicted the exact peptidoform for 69.2% of the spectra in a public phosphopeptide-enriched dataset and for 66 to 72% of the phosphopeptide spectra in three phosphoproteomic studies from our laboratory that were not used for training. In these studies, it identified numbers of phosphopeptides comparable to or larger than those of database search engines, and proportionally more where database search identified fewer. For the phosphopeptides identified only by Casanovo-TMT, entrapment estimated that more than 90% were correct, and an independent search engine assigned the same sequence to 95 to 98% of the spectra that it identified with confidence. Casanovo-TMT can therefore extend database search in TMTpro-based quantitative proteomics, particularly in studies in which database search identifies few phosphopeptides.

## INTRODUCTION

Isobaric labeling with tandem mass tags is the principal route to multiplexed quantitative proteomics [1, 2]. The current reagent, TMTpro, is supplied in a 16-plex and an 18-plex set whose members are isobaric. Every channel of either set adds the same 304.2071 Da to a peptide, and the sets differ only in where the heavy isotopes sit, which is read from the reporter ions. Sixteen or eighteen samples are labeled, pooled and acquired together, and abundance is read from those ions [3]. The reagent acylates the peptide N terminus and every lysine side chain, adding 304.2071 Da at each position [4, 5]. Labeled and unlabeled spectra therefore differ in precursor mass and across the whole b- and y-ion series, not only in the low-mass reporter region. Peptides in these experiments are almost always assigned by peptide spectrum matching via database search, with the tag declared as a fixed modification [6]. A spectrum whose peptide is not in that database cannot be assigned, however well it fragmented.

De novo sequencing derives a peptide sequence from the fragment ions alone, without requiring the peptide to be present in a database [7, 8]. Deep learning has made de novo sequencing practical at proteome scale. Models such as Casanovo and InstaNovo, trained on millions of peptides identified by database search, now perform nearly as well as database search itself, at least for unmodified tryptic peptides [9, 10]. Casanovo, the model we build on, is a transformer that emits one residue at a time and explores alternatives by beam search, which carries a fixed number of partial sequences forward at each step [9]. It was pretrained on MassIVE-KB, a community spectral library assembled by database search of public human datasets [11], and no isobaric tag was among the modifications searched to build it. The tokens Casanovo can emit, its residue vocabulary, are fixed before training and comprise the twenty amino acids plus carbamidomethylated cysteine, oxidized methionine, deamidation and a small set of N-terminal groups [9, 12]. A modification absent from the vocabulary when training began cannot appear in any prediction, whatever the spectrum contains.

Several studies have extended existing de novo models to modifications absent from their vocabularies [13–15]. A combination of an extended vocabulary and end-to-end fine-tuning has been applied to phosphorylation [14, 15] and, more broadly, to twelve residue-modification combinations absent from the base vocabulary, including lysine acetylation, ubiquitination, methylation and O-linked glycosylation [13]. Moreover, version 5.2 of Casanovo now provides a configuration option that adds tokens to a pretrained checkpoint and initializes each of them from an existing residue [12]. Publicly released trained models now cover phosphorylation [14] and the twelve combinations above [13], yet none appears to cover an isobaric label such as TMTpro. We inspected the residue vocabularies distributed with the current de novo models and found no released checkpoint able to emit a TMTpro residue. None declares the 304.2071 Da increment, including the extended vocabulary file of 84 modification tokens distributed with the InstaNovo package [9, 10, 13–15]. PROSPECT PTMs, a large reference corpus of modified synthetic peptides, includes TMT-labeled subsets, but these carry the earlier 229.1629 Da reagent, and its authors state that it contains no TMTpro-labeled peptides [16]. Against this background, the limiting factor appears to be the training data rather than the architecture, since fine-tuning needs labeled spectra with trustworthy sequence assignments in the millions. Conventionally, TMTpro-based proteomics relies almost entirely on database search, and the spectra it cannot assign, from sequence variants, unannotated peptides or unexpected modifications, have no other route to a sequence. Yet de novo sequencing of such spectra is increasingly in demand, and a deep learning model for TMTpro-labeled peptides would address an unmet need.

In this study, we fine-tuned Casanovo on a corpus of 21.0 million in-house Orbitrap spectra, combining TMTpro-labeled and label-free spectra, and evaluated the resulting model (Fig. 1). In addition, the labeled spectra included 2.4 million phosphopeptide spectra (13.0%) from phosphopeptide-enriched runs. We represented the tag as explicit tokens at the peptide N terminus and on lysine, using the extended-vocabulary procedure of Casanovo rather than a new architecture [12]. For decoding we applied a knapsack-style rule, adapted from DeepNovo [17], that re-ranks the candidates held by the beam against the measured precursor mass [18]. We refer to the resulting model as Casanovo-TMT and evaluated it on spectra drawn from the test partition of the corpus, on public datasets from other laboratories, one of them phosphopeptide-enriched, and on three phosphoproteomic studies from our laboratory that were not used in training. On the TMTpro-labeled test spectra, Casanovo-TMT sequenced 84.7% of peptidoforms exactly, whereas the released Casanovo, which lacks a TMTpro token, sequenced 0.3%. On two public global proteomic datasets, precision was 87.4 and 79.6%. On a public phosphoproteomic dataset, precision was 69.2%. In those three studies, Casanovo-TMT identified 19.7 to 44.8% more phosphopeptides than SEQUEST HT. We make Casanovo-TMT openly available, both as trained weights and as a desktop application that requires no programming (https://github.com/existstar4-gif/casanovo-tmtpro), so that any laboratory using TMTpro can apply de novo sequencing to its own data.

**Figure 1.**
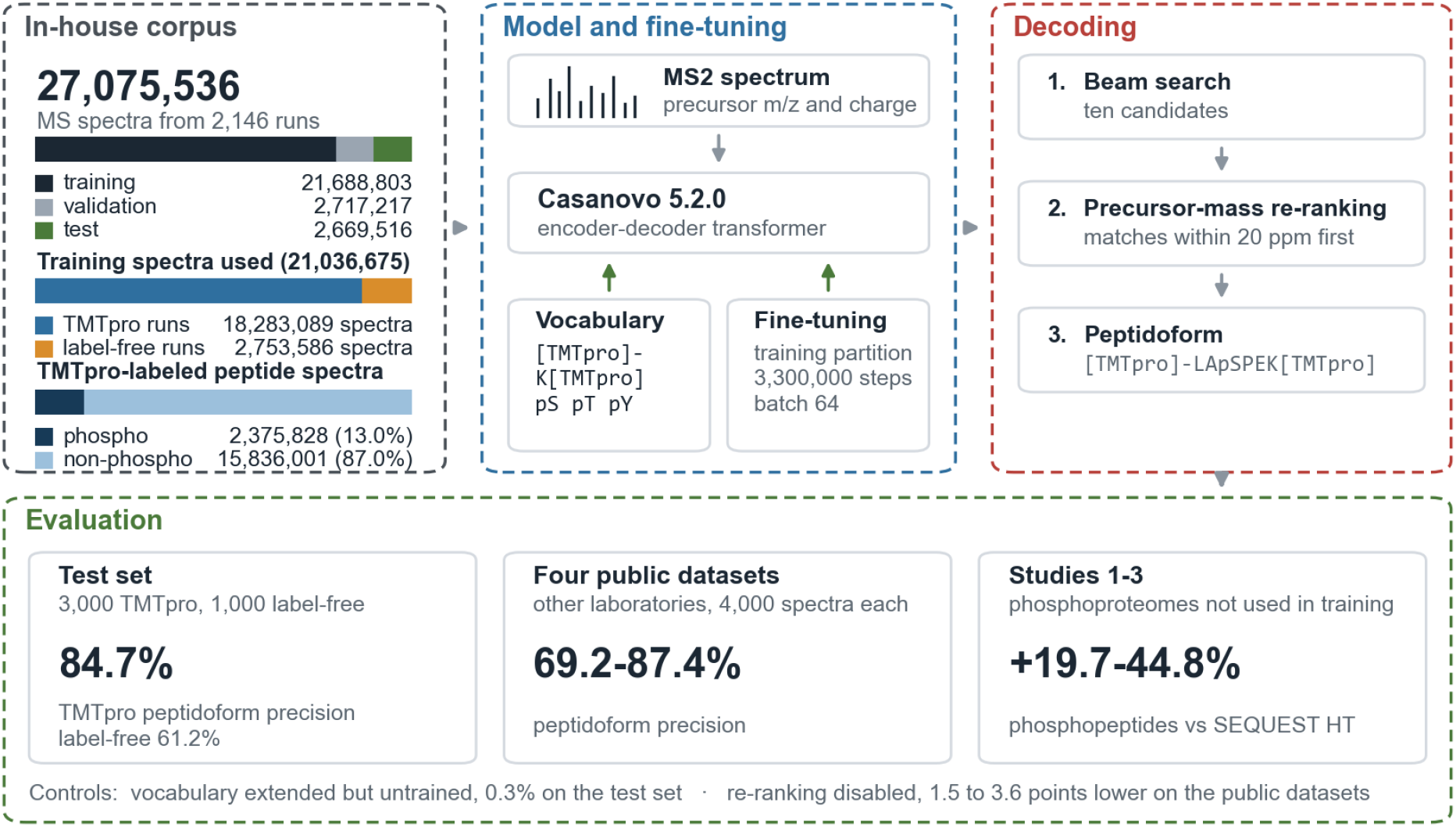
Corpus, model, decoding and evaluation. The corpus comprises 27,075,536 MS spectra from 2,146 runs acquired in our own laboratory, divided by peptide into training, validation and test partitions so that a peptide falls entirely in one partition; the counts of spectra from TMTpro and label-free runs, and the numbers of phosphorylated and non-phosphorylated TMTpro-labeled peptide spectra, are those of the 21,036,675 training spectra used for fine-tuning. Casanovo 5.2.0 was fine-tuned on the training partition after its residue vocabulary was extended with the two TMTpro tokens and the phosphorylation tokens. At inference the beam retains ten candidates, and candidates whose peptidoform mass matches the measured precursor within 20 ppm are ranked above the others; none is discarded, so every spectrum still receives a prediction. Evaluation used a test set of 4,000 spectra drawn from the test partition, four datasets acquired in other laboratories, and three phosphoproteomic studies from our laboratory that contributed no spectra to training.

## EXPERIMENTAL PROCEDURES

### Corpus Assembly and Mass Verification

All spectra in the corpus were acquired in our laboratory, in 2,146 LC-MS/MS runs on an Orbitrap Exploris 480 mass spectrometer (Thermo Fisher Scientific, Waltham, MA). None of these runs had been deposited in a public repository before this work, so the training corpus does not overlap any published benchmark. Peptidoforms were assigned to the spectra by database search with SEQUEST HT in Proteome Discoverer 3.3 (Thermo Fisher Scientific) against the UniProtKB human or mouse proteome (release 2025_01), chosen by the species of the sample, and only rank-1 peptide-spectrum matches of high confidence with a q-value below 0.01 were retained.

The corpus comprised 27,075,536 spectra. They were partitioned by peptide rather than by spectrum or by run: the unmodified sequence of each peptidoform, with isoleucine and leucine equated, was hashed and assigned to training, validation or test in proportions of 80, 10 and 10, so that a peptide falls in the same partition wherever it occurs. The partitions held 21,688,803, 2,717,217 and 2,669,516 spectra, respectively; when the training partition was loaded for fine-tuning, the 652,128 spectra with a precursor charge above 4 or fewer than 20 peaks were excluded, leaving 21,036,675. The test set was drawn from the 2,669,516 spectra of the test partition, none of which was used in training.

Mass consistency of the assignments was audited across all 27,075,536 spectra of the corpus by comparing the theoretical neutral mass of each assigned peptidoform with the mass implied by the recorded precursor m/z and charge; 99.94% agreed within 10 ppm and 55.9% within 1 ppm. The 15,551 spectra (0.057%) that disagreed by more than 50 ppm all came from TMTpro runs, and 95.6% of them exceeded the mass of the assigned peptidoform by 42.011 Da, that of an unassigned acetyl group; they were retained, since at that rate they cannot materially affect fine-tuning.

Of the 21,036,675 spectra used for fine-tuning, 2,753,586 (13.1%) came from label-free runs and the remainder from TMTpro runs. Classified by the peptide rather than by the run, 18,211,829 spectra had a TMTpro-labeled reference, 2,375,828 (13.0%) of phosphorylated and 15,836,001 (87.0%) of non-phosphorylated peptides; this count is 71,260 lower than that of spectra from TMTpro runs because a peptide with an acetylated N terminus and no lysine carries no tag. Across the corpus, 3,082,088 spectra had a phosphorylated reference, and of the 3,639,785 phosphosites they carried, 86.0% were on serine, 12.5% on threonine and 1.5% on tyrosine. This composition reflects the sample preparation: phosphopeptides were enriched manually on Fe(III)-nitrilotriacetic acid resin, which binds the phosphate group regardless of the residue, and no anti-phosphotyrosine enrichment was performed.

The TMTpro-labeled runs used the 16-plex reagent set. The 135N reporter, which only the 18-plex set carries and which has no neighbor within 1 Da, was present in 0.6% of the test spectra, a rate consistent with coincidental peaks; the other channel added by the 18-plex set lies 6.3 mDa from a 16-plex channel and cannot be distinguished at the tolerance used here.

### Model, Residue Vocabulary and Fine-Tuning

Fine-tuning started from the released Casanovo checkpoint [9, 12], a transformer with 9 layers, a model dimension of 512, 8 attention heads, a feed-forward dimension of 1,024 and no dropout; the maximum precursor charge of 4 is fixed by the charge embedding of this checkpoint. Two TMTpro tokens were added to the residue vocabulary, an N-terminal [TMTpro]-of 304.207146 Da and K[TMTpro] of 432.302109 Da, together with tokens for phosphoserine, phosphothreonine and phosphotyrosine. Each new output-layer row was initialized by copying the row of a related token already in the model: [Acetyl]-for the N-terminal tag, unmodified lysine for the labeled lysine and the unmodified residue for each phosphorylation token. Tokens for phosphoarginine and phospholysine were also added, only so that the spectra whose references carry these sites could be read; these 40 sites are 0.001% of the phosphosites in the corpus, and we regard them as localization artifacts rather than as a target of the model.

Fine-tuning used the Adam optimizer with a batch size of 64, a learning rate of 1.5e-4 with 10,000 warm-up iterations and a cosine schedule of period 3,310,000, weight decay of 1.0e-5 and label smoothing of 0.01. Spectra were preprocessed by retaining at most the 150 most intense peaks above a relative intensity of 0.01 and at least 20 peaks, within 50 to 2,500 m/z, with peaks within 2.0 m/z of the precursor removed, and peptides shorter than 6 or longer than 100 residues were excluded. Training ran for 3,300,000 steps, about 10 passes over the training partition, in about 12.5 days of computation on a single NVIDIA GeForce RTX 5070 Ti, with validation every 5,000 steps, and the checkpoint at the last step was used for every result reported here.

The training objective was the token cross-entropy with an auxiliary hinge term. Because Casanovo decodes from the C terminus, the N-terminal tag is the last residue token emitted, in direct competition with the stop token. The term acts only at positions whose target is the N-terminal [TMTpro]-token, where it penalizes a difference of less than 5 between the logit of that token and that of the stop token, with a weight of 1.0; other positions are unaffected, and the validation loss excludes the term. The term was active throughout training. In a paired ablation at 40,000 steps, precision with and without the term differed by 0.0 percentage points (paired 95% CI ±1.2 points), but one branch of that ablation was restarted twice, so the ablation is inconclusive and the contribution of the term was not established.

The order in which training spectra were read was changed once during training. For steps 0 to 2,360,000 they were read in file-name order, in which each study forms a contiguous block of a single class and the label-free spectra precede the TMTpro spectra in every pass; because a shuffle buffer of 5,000 spectra cannot mix blocks of that size, the training partition was rewritten with TMTpro and label-free spectra interleaved, and steps 2,360,000 to 3,300,000 read the interleaved copy. Training was not repeated from the start with the interleaved order.

### Decoding

Predictions were generated with a beam width of 10, with the precursor mass tolerance of the Casanovo configuration set to 50 ppm and isotope errors of 0 to 1. Casanovo 5.2.0 does not constrain de novo output by the measured precursor mass, so we restored this constraint when ranking the candidates held in the beam: candidates whose peptidoform mass matched the measured precursor within 20 ppm, allowing one carbon-13 isotope, were ranked above the others, and when no candidate matched, the highest-scoring candidate was reported. No candidate is discarded, so every spectrum receives a prediction with and without the constraint, and the two settings can be compared spectrum by spectrum. DeepNovo imposed the same constraint during generation, excluding at each step the residues whose mass could not complete the remaining precursor mass [17]; applying it after generation cannot recover a peptide the beam never produced, but it leaves generation unchanged.

Candidates of equal mass, such as sequences that differ only in the order of two residues, cannot be separated by the precursor mass, but only one of them may occur among the peptides of a tryptic digest of the reference proteome, which we refer to as the tryptic search space. Where this second stage was applied, candidates present in the tryptic search space were ranked first within each of the two mass groups, so that a candidate whose mass contradicts the precursor cannot be ranked above one that matches it because it occurs in the database. The tryptic search space was built from the reference proteome of the species, allowing up to two missed cleavages and peptide lengths of 6 to 50, with isoleucine and leucine equated, and included the peptides that begin at the second residue of proteins whose first residue is methionine.

### Evaluation

Peptidoform precision was defined as the percentage of evaluated spectra whose prediction matched the reference peptidoform exactly, including every modification, with isoleucine and leucine treated as identical; spectra without a prediction were counted as incorrect. This criterion is stricter than the peptide precision reported for Casanovo and DeepNovo [9, 17], which accepts a residue whose mass differs from the reference by less than 0.1 Da and a prefix whose cumulative mass differs by less than 0.5 Da, and therefore counts substitutions such as glutamine for lysine as matches; the values reported here are not comparable with published peptide precisions. Predictions were matched to references by raw file and scan number.

Confidence intervals are 95% cluster bootstrap intervals from 10,000 resamples in which peptides rather than spectra were drawn with replacement, all spectra of a drawn peptide being included together. The 4,000 spectra of the test set represent 823 distinct peptides, the most frequent of which accounts for 4.3% of the spectra, so intervals that treat spectra as independent would understate the uncertainty. Spectra from the same study may also be correlated, but intervals that resample studies could not be computed because the study of origin of each test spectrum was not recorded; in an earlier analysis such intervals were about 1.7 times wider than peptide-level intervals, so the intervals reported here may understate the uncertainty.

Datasets that contributed no spectra to training are called independent datasets throughout: the four public datasets retrieved from PRIDE described below and three phosphoproteomic studies from our laboratory that were not used for training. The test set is kept distinct from them: its spectra come from the same acquisitions as the training spectra and are separated from them by peptide sequence, whereas the independent datasets come from different samples.

Public datasets were retrieved from PRIDE. TMTpro-labeled datasets were required to supply both peak lists and the depositors’ search results and to retain the reporter-ion region between 126 and 135 m/z, and three met these requirements; a label-free dataset that supplied both peak lists and search results was added to evaluate unlabeled peptides. From each dataset, 4,000 spectra with a reference identification were selected at random. The four datasets are described in Supplementary Table S1.

Three phosphoproteomic studies from our laboratory served as independent datasets for the comparison with database search: Studies 1 and 2, each in 12 fractions labeled with 18-plex TMTpro, and Study 3, in 24 fractions labeled with 16-plex TMTpro, all from mouse tissue. None of their 49 raw files was among the source files of the training corpus; one fraction of Study 1 had been acquired twice, and both acquisitions were included, as in the database search. The raw files had been searched in Proteome Discoverer [19] with SEQUEST HT [20] and, separately, with CHIMERYS [21], each at a 1% false discovery rate. Peptidoform precision on these studies was computed as defined above against the high-confidence phosphopeptide-spectrum matches of SEQUEST HT. For Studies 1 to 3, the candidates in the beam were additionally re-ranked by the tryptic search space of the mouse reference proteome (Fig. 2B).

**Figure 2.**
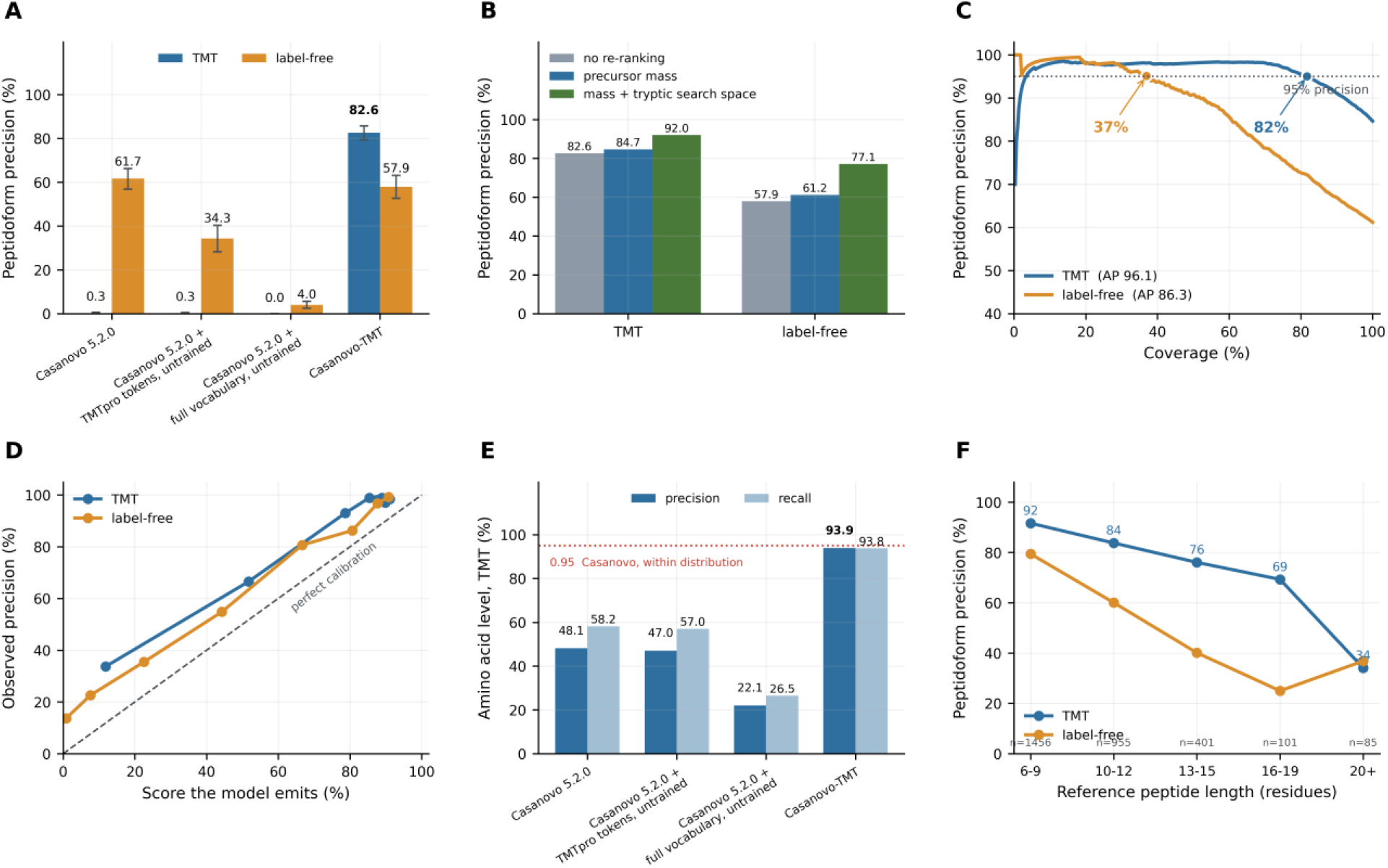
Performance of Casanovo-TMT on the test set. The test set comprises 3,000 TMTpro-labeled spectra (TMT) and 1,000 spectra of unlabeled peptides (label-free) selected from the test partition. Peptidoform precision is the percentage of evaluated spectra whose predicted peptidoform, including every modification and TMTpro tag, matched the reference peptidoform exactly, with isoleucine and leucine treated as identical; spectra for which no peptide was predicted are counted as incorrect. All conditions were decoded with a beam of ten candidates. *A*, Peptidoform precision under four conditions decoded with identical settings, all without precursor-mass re-ranking: Casanovo 5.2.0, the released weights; Casanovo 5.2.0 + TMTpro tokens, untrained, the released weights with the two TMTpro tokens, [TMTpro]- and K[TMTpro], added to the vocabulary and no further training; Casanovo 5.2.0 + full vocabulary, untrained, the released weights with the two TMTpro tokens and five phosphorylation tokens added and no further training; and Casanovo-TMT. Error bars are 95% confidence intervals from 10,000 bootstrap resamples in which peptides rather than spectra were drawn with replacement, all spectra of a drawn peptide being included together. *B*, Peptidoform precision of Casanovo-TMT under three ways of ordering the ten candidates held in the beam: no re-ranking; precursor mass, in which candidates whose peptidoform mass matches the measured precursor within 20 ppm, allowing one carbon-13 isotope, are ranked above the others; and mass + tryptic search space, in which candidates present in a tryptic digest of the reference proteome of the species are additionally ranked first within each of the two mass groups. No candidate is discarded, so the same spectra are scored in all three. Precursor mass is the setting used for Casanovo-TMT in *C*, *D* and *F* and, unless stated otherwise, throughout this work. *C*, Coverage-precision curves of Casanovo-TMT. Predictions were ranked by the score the model assigns to each, and a point at coverage *x* gives the peptidoform precision of the *x*% of spectra with the highest scores; a coverage of 100% corresponds to the precursor-mass values in *B*. The dotted line marks 95% precision, and circles mark the largest coverage at which precision remains at or above it, with that coverage printed beside each circle. AP, average precision, the area under the curve, in percent. *D*, Calibration of the score of Casanovo-TMT. Predictions were sorted by score and divided into eight bins of equal size; each point plots the mean score of a bin, expressed as a percentage, against the peptidoform precision observed in that bin. The dashed diagonal marks a score that equals the observed precision. *E*, Amino acid-level precision and recall on the TMT spectra under the four conditions of *A*, all without precursor-mass re-ranking, scored with the mass-tolerant criterion described previously [9, 17]. A predicted residue is counted as correct when its mass agrees with the reference residue within 0.1 Da and the cumulative mass of the residues before it agrees within 0.5 Da; precision is the percentage of predicted residues that are correct and recall the percentage of reference residues that are recovered. The dotted line marks the amino acid-level precision of 0.95 reported for Casanovo on data from its own training distribution. These values count residues rather than whole peptidoforms and are not comparable with the peptidoform precision of the other panels. *F*, Peptidoform precision of Casanovo-TMT against the length of the reference peptide, in residues; n is the number of TMT spectra in each length bin.

For counting, peptides were compared as bare sequences with isoleucine and leucine equated, and a sequence was counted as a phosphopeptide if it had been observed with a phosphate. Predictions of Casanovo-TMT were thresholded by target-decoy competition against a decoy digest of the reversed protein sequences, generated with the same digestion rules as the tryptic search space: the threshold was the score at which matches to the decoy digest reached 1% of matches to the target digest, computed separately for phosphorylated and non-phosphorylated sequences as described below, and sequences outside the protein database were excluded. Because this threshold estimates how often a sequence falls in the tryptic search space by chance rather than whether it is correct, it is not equivalent to the false discovery rate of a search engine. As a second search engine, Sage 0.14.6 [22] was run on the same spectra against the Swiss-Prot mouse reference proteome of 17,229 entries, with TMTpro on peptide N-termini and lysine and carbamidomethylation of cysteine as fixed modifications, phosphorylation of serine, threonine and tyrosine and oxidation of methionine as variable modifications with at most two per peptide, two missed cleavages, tolerances of 10 ppm for precursors and 20 ppm for fragments, and peptides of 6 to 40 residues and up to 5,000 Da, limits set by the memory available for the fragment index.

Three rules made the counts comparable across methods. First, every method was assessed against the protein database used by the Proteome Discoverer searches, the Swiss-Prot mouse reference proteome of 17,229 entries together with 4,604 unreviewed entries for genes absent from it. Second, predictions of Casanovo-TMT without an N-terminal TMTpro tag were removed before any threshold was computed, because the database searches fixed the tag on peptide N termini and cannot return an unlabeled peptide; protein N-terminal acetylation was retained. Third, the decoy threshold of Casanovo-TMT was computed separately for phosphorylated and non-phosphorylated sequences, each at 1%, and the two thresholds were applied together, giving a combined rate of 0.98 to 1.00%. FragPipe 22 with MSFragger and Percolator [23–25] was run on the same raw files with its TMT16 phosphopeptide workflow and the same database, and its default output, filtered by Philosopher at a 1% false discovery rate at the peptide-spectrum match, peptide and protein levels, was used; SEQUEST HT and CHIMERYS were likewise used at their own 1% thresholds, as the high-confidence matches exported by Proteome Discoverer. A phosphopeptide was counted as quantifiable when at least one of its spectra carried a complete TMTpro label, contributed at least half of the precursor signal in its isolation window and had reporter intensity in at least eight channels, and this rule was applied to the spectra of every method in the same way.

Phosphopeptides identified by Casanovo-TMT and by at least one database search engine (shared) or by none (unique) were each represented by their highest-scoring spectrum. False discovery proportions were estimated by entrapment with a tryptic digest of the *Arabidopsis thaliana* Swiss-Prot proteome from which peptides present in the mouse target or decoy digests were removed (1,721,553 peptides, 0.667 times the mouse digest), using the combined estimator [26]; this digest did not enter the threshold calculation. Agreement with Sage was the fraction of these spectra that Sage, at a 1% spectrum-level false discovery rate, assigned to the same sequence. Retention time was predicted within each fraction by gradient boosting in scikit-learn, trained on the shared phosphopeptides with five-fold out-of-fold prediction. Reporter-ion patterns, log2-transformed and normalized by channel, were compared by Pearson correlation with the median pattern of database-confirmed phosphopeptides carrying the same phosphosite, since the abundance of a phosphosite need not follow that of its protein.

### Software and Reproducibility

Analyses used Python 3.12.13, PyTorch 2.11.0 with CUDA 12.8, Casanovo 5.2.0, depthcharge-ms 0.4.10, Lightning 2.6.5 and scikit-learn 1.9.0 on Windows 11, and raw files were converted to mzML with ThermoRawFileParser 2.0.0. Three files of the installed Casanovo distribution were modified: denovo/model.py (254 lines added, 7 removed), denovo/model_runner.py (9 added) and data/ms_io.py (26 added, 1 removed); the modified files and their differences from the released 5.2.0 source are provided with the code, since the results cannot be reproduced from the package as distributed. One of these changes works around a defect in beam search that discards every beam carrying an N-terminal modification, so that an N-terminally labeled peptide cannot be predicted by the unmodified 5.2.0 release; the defect is fixed upstream in 5.2.1 [18]. Database searches used SEQUEST HT and CHIMERYS in Proteome Discoverer 3.3, FragPipe 22 with MSFragger 4.1, Percolator 3.06.5 and Philosopher 5.1.1, and Sage 0.14.6; the public datasets were searched for comparison of reference identifications with FragPipe 24 with MSFragger 4.4.1, Percolator 3.7.1 and Philosopher 5.1.3.

### Experimental Design and Statistical Rationale

The design comprised one training corpus and three kinds of evaluation data. The corpus of 27,075,536 spectra from 2,146 runs was partitioned by peptide into training, validation and test partitions in proportions of 80, 10 and 10, so that no peptide occurred in more than one partition; the validation partition was used only to monitor training. Performance was evaluated on a test set of 4,000 spectra from the test partition, 3,000 from TMTpro-labeled and 1,000 from label-free runs; on 4,000 randomly selected spectra from each of four public datasets that contributed no spectra to training; and on all 49 raw files of three phosphoproteomic studies from our laboratory that were not used for training. Biological replicates were not part of the design, because the unit of evaluation is a spectrum and its reference identification rather than a biological condition, and each study was analyzed once by each method. Controls were the released Casanovo weights and the same weights with the extended vocabulary but without training (Fig. 2A), Casanovo-TMT without precursor-mass re-ranking (Fig. 2B) and, for the identification of phosphopeptides, three database search engines applied to the same raw files (Fig. 4). Precision is reported with 95% confidence intervals from 10,000 bootstrap resamples in which peptides rather than spectra were drawn, because spectra of the same peptide are not independent. The relation between the gain of Casanovo-TMT over SEQUEST HT and the number of phosphopeptides identified by SEQUEST HT was tested by Spearman rank correlation over seven studies with an exact two-sided P value. Identifications of Casanovo-TMT were thresholded by target-decoy competition at 1%, and the error among the phosphopeptides identified by Casanovo-TMT alone was estimated by entrapment with a foreign-species digest [26]. Sample sizes were not determined by power analysis; with the test set of 4,000 spectra, the 95% confidence interval of precision on TMTpro-labeled spectra spanned 81.6 to 87.5%.

### De-identification

Sample identifiers in the released corpus carry study, cell-line, gene and specimen names in free text. Every such identifier was replaced by an opaque token before release, using a mapping of 2,146 sample identifiers that is retained by the authors and is not distributed. The 164 tokens replaced comprise gene and protein names, cell-line designations, disease-model names, specimen types and project names. Peptide sequences themselves are released unchanged, since they are tryptic products of reference proteomes and do not encode subject-specific variation.

## RESULTS

### Design of the Corpus, the Model and the Evaluation

We assembled a corpus of 27,075,536 spectra from 2,146 Orbitrap runs acquired in our own laboratory and not previously deposited, of which 1,838 (85.6%) are TMTpro-tagged runs and 308 (14.4%) label-free runs. Of the 21,036,675 spectra used for fine-tuning, 18,211,829 had a TMTpro-labeled reference, 2,375,828 (13.0%) of phosphorylated and 15,836,001 (87.0%) of non-phosphorylated peptides. The phosphopeptides were enriched by immobilized metal affinity chromatography on Fe(III)-nitrilotriacetic acid resin, and the phosphosites carried by the phosphopeptides of the corpus are accordingly 86.0% phosphoserine, 12.5% phosphothreonine and 1.5% phosphotyrosine. We split the corpus by peptide into training (21,688,803 spectra), validation (2,717,217) and test (2,669,516) partitions, so that every form of a peptide, labeled or unlabeled and with or without phosphorylation, falls in one partition. We then fine-tuned Casanovo 5.2.0 end to end on the training partition for 3,300,000 steps at a batch size of 64, with its residue vocabulary extended; we refer to the resulting model as Casanovo-TMT. Because Casanovo emits the N-terminal tag last, in competition with the stop token, an auxiliary hinge term was added to the token cross-entropy during training; it acts only where the reference peptide carries the tag, favoring the tag over the stop token at that position, and plays no part in inference. At inference, the beam retains the ten highest-scoring candidate peptides for each spectrum. Candidates whose mass matches the measured precursor mass are ranked above those that do not, and when no candidate matches, the highest-scoring candidate is reported, so every spectrum still receives a prediction. Casanovo-TMT was evaluated on a test set of 4,000 spectra drawn from the test partition, whose spectra were not used in training, on four datasets acquired in other laboratories, and on three phosphoproteomic studies from our laboratory (Fig. 1).

### Fine-Tuning of Casanovo With an Extended TMTpro Residue Vocabulary

To let the decoder emit the TMTpro tag as part of the sequence itself, we extended the residue vocabulary, the set of symbols a de novo model may write at each position. Two tokens were added: a peptide N-terminal [TMTpro]-of 304.207146 Da and a lysine K[TMTpro] of 432.302109 Da. Each was initialized by copying the output-layer row of a symbol already present in the model, [Acetyl]- and unmodified K, respectively. Tokens for phosphoserine, phosphothreonine, and phosphotyrosine were added on the same basis, each from its unmodified residue. Two further tokens, phosphoarginine and phospholysine, were retained only so that the spectra whose references carry these sites could be read; these 40 sites account for 0.001% of the phosphosites in the corpus.

### Performance of Casanovo-TMT on the Test Set

We evaluated Casanovo-TMT on 4,000 spectra selected from the test partition (Fig. 1). A prediction was counted as correct only when the complete peptidoform, tag included, matched the reference peptidoform assigned to that spectrum by database search, with leucine and isoleucine treated as identical because they are isobaric. Peptidoform precision is the number of correct predictions divided by the number of spectra evaluated; spectra for which the model returned no prediction remain in the denominator and count as incorrect. At 100% coverage, that is, with every prediction retained whatever score the model assigned to it, precision was 82.6% on the 3,000 TMT spectra (95% CI 79.3–85.7) and 57.9% on the 1,000 label-free spectra (52.6– 63.1), and 76.5% over the whole set (73.4–79.3) (Fig. 2A). Confidence intervals were obtained by resampling peptides.

To separate the contribution of the added tokens from that of training on labeled spectra, we sequenced the same test set with three further configurations: the released Casanovo weights; the released weights with only the two TMTpro tokens added; and the released weights with the full extended vocabulary described above, which also includes the five phosphorylation tokens. In both configurations, the added tokens were copy-initialized as described above, and no further training was performed (Fig. 2A). We decoded all of them with a beam width of 10, meaning that ten partial sequences were carried forward at each decoding step, and scored all against the same reference peptidoforms. The released Casanovo, whose vocabulary contains no TMTpro token, reached 0.3% on TMT spectra and 61.7% on label-free spectra. Copy-initializing the two TMTpro tokens and withholding all training left TMT precision at 0.3%, while label-free precision fell to 34.3% (Fig. 2A). With the full extended vocabulary, label-free precision fell further to 4.0%, apparently because each copied token received the same score as the token it was copied from, so that the choice between them was arbitrary (Fig. 2A). The vocabulary-extended but untrained model gained nothing on TMT spectra, indicating that the 82.6% is a product of training on labeled spectra and not of the change to the vocabulary.

In de novo mode, Casanovo 5.2.0 no longer uses the measured precursor mass to constrain the output. We restored that constraint as the re-ranking step described above rather than as a filter. Because no candidate was removed, every spectrum received a prediction in both decoding modes, so the two modes could be compared on identical spectra. On the test set the constraint raised precision from 82.6 to 84.7% on TMT spectra and from 57.9 to 61.2% on label-free spectra (Fig. 2B). With re-ranking, which is used for Casanovo-TMT throughout the rest of this work, precision on the test set was 84.7% on TMT spectra (95% CI 81.6–87.5), 61.2% on label-free spectra (56.2–66.0) and 78.8% overall (76.0–81.4).

When a reference proteome is available, re-ranking can use it as a second criterion: among candidates that agree equally with the precursor mass, those present in an in silico tryptic digest of the proteome, the tryptic search space, are ranked first (Experimental Procedures). With this second stage, precision on the test set rose from 84.7 to 92.0% on TMT spectra and from 61.2 to 77.1% on label-free spectra (Fig. 2B). Casanovo-TMT can therefore also be used in a database-assisted mode.

In addition, we examined peptidoform precision at each coverage, the percentage of spectra retained when the predictions are ranked by the score Casanovo-TMT assigns to each and only the highest-scoring are kept (Fig. 2C). On the TMTpro spectra of the test set, precision was 95% at a coverage of 81.7%, 90% at a coverage of 92.2%, and 84.7% at a coverage of 100%. On the label-free spectra, precision was 95% at a coverage of 37.0%, 90% at a coverage of 51.8%, and 61.2% at a coverage of 100%. Thus, Casanovo-TMT reached high precision once its lowest-scoring predictions were excluded.

We next examined whether the score of Casanovo-TMT can be read as the probability that a prediction is correct (Fig. 2D). Predictions on TMTpro and on label-free spectra were sorted separately by score and divided into eight bins of nearly equal size, and the mean score of each bin was compared with the precision observed in it. In every bin, the observed precision exceeded the mean score; for example, the 371 TMTpro predictions in the bin with a mean score of 0.854 were correct 98.9% of the time. The score therefore ranks predictions reliably but underestimates how often they are correct, so a threshold should be chosen from the coverage-precision curve rather than from the score itself.

We next compared Casanovo-TMT with published de novo models, which report amino acid-level precision and recall rather than peptidoform precision (Fig. 2E). For this comparison, the same predictions were scored with the mass-tolerant criterion described previously [9, 17]. Without re-ranking, as for the other conditions in Fig. 2E, amino acid-level precision on the TMTpro spectra was 93.9% and recall 93.8%; with re-ranking, they were 94.7% and 94.6%. These values match those that the developers of Casanovo and DeepNovo report on data from their own training distribution, although here they were reached on spectra carrying a modification absent from their vocabularies.

We then examined how precision depends on peptide length (Fig. 2F). Spectra were grouped by the length of the reference peptide, and peptidoform precision was computed in each group. On the TMTpro spectra, precision fell from 91.6% at six to nine residues to 34.1% at twenty or more residues, and the label-free spectra showed the same decline. Long peptides are therefore the weakest population for Casanovo-TMT, as they are for de novo sequencing generally. Altogether, these results indicate that extending the residue vocabulary of Casanovo with the TMTpro tag and fine-tuning it on TMTpro-labeled spectra yields a model that sequences TMTpro-labeled peptides with high precision, and whose score effectively separates correct from incorrect predictions.

### Precision Across Training Steps

To examine how precision depends on the amount of training, we evaluated five checkpoints of a single training run on the test set under identical decoding with precursor-mass re-ranking, at 20,000, 40,000, 80,000, 160,000 and 320,000 training steps. Overall precision rose from 75.8 to 77.8% across that sixteenfold increase, TMTpro precision from 81.2 to 83.2% and label-free precision from 59.9 to 61.4%. The rate at which a TMTpro token was emitted on a label-free spectrum was 0.0% at every point. Casanovo-TMT, trained for 3,300,000 steps, reached 84.7% on the TMTpro spectra of the same set. Precision within the longest length stratum moved erratically between adjacent checkpoints, between 13.5 and 40.4% on 104 spectra, and no claim about length generalization is made from these data. Most of the gain in precision was therefore reached within the first 20,000 steps, and further training added only a few percentage points.

### Validation of Peptidoform Precision on Four Independent Public Datasets

To determine whether the precision measured on our own data is reproduced on data from other laboratories, we analyzed public datasets that were not used in training (Fig. 3A). The four datasets are described in Supplementary Table S1. Peptides were sequenced with the same settings as the test set, with precursor-mass re-ranking but without the tryptic search-space stage. Predictions were scored against the peptide identifications that the depositors obtained by database search. From each dataset, 4,000 spectra with a reference identification were selected at random.

**Figure 3.**
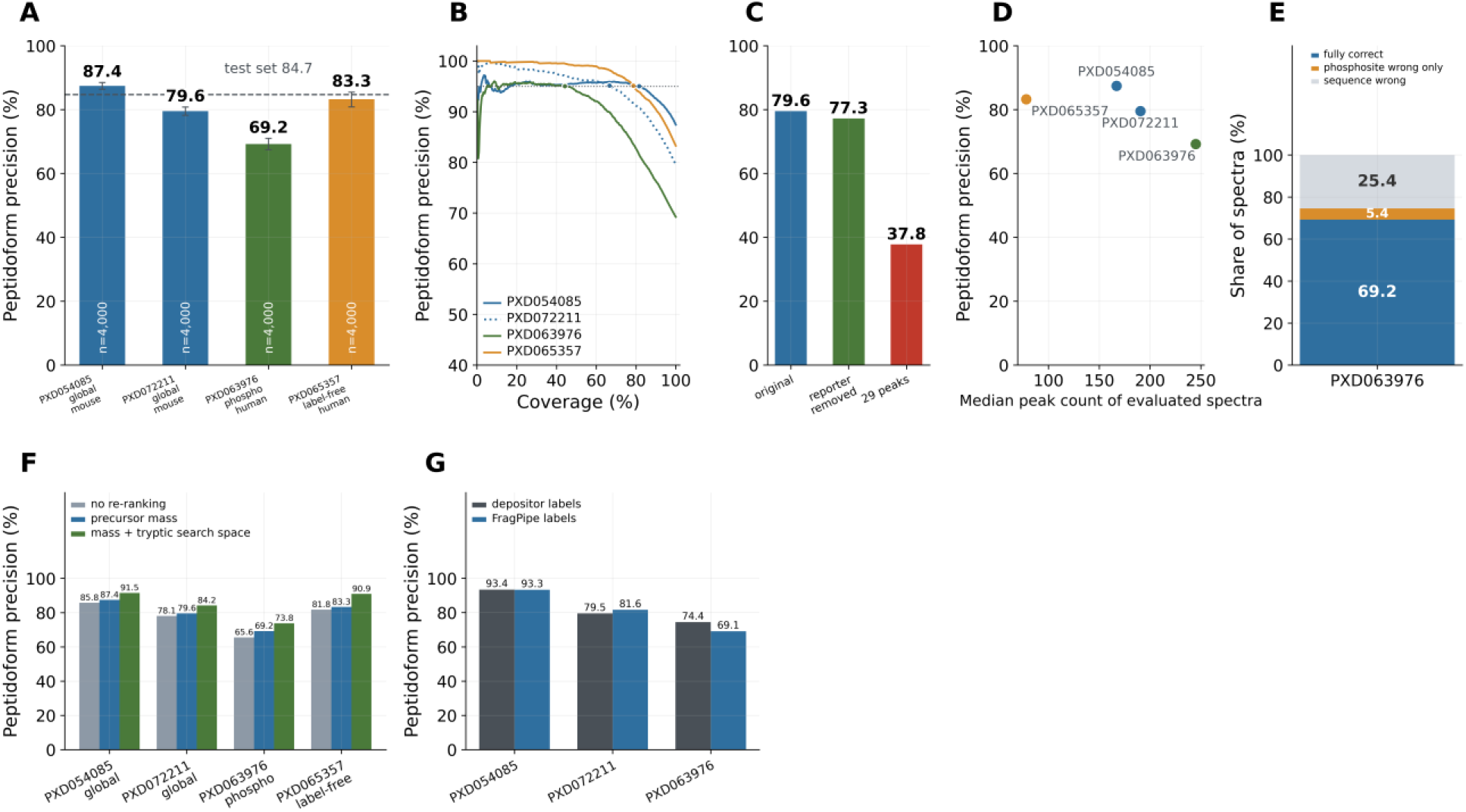
Performance of Casanovo-TMT on four public datasets. The four datasets contributed no spectrum to training and are described in Supplementary Table S1: PXD054085 and PXD072211, TMTpro-labeled mouse global-proteome datasets; PXD063976, a TMTpro-labeled human phosphopeptide-enriched dataset; and PXD065357, a human label-free dataset. From each dataset, 4,000 spectra with a reference identification were selected at random, and predictions were scored against the peptide identifications that the depositors obtained by database search. Peptidoform precision is defined as in Fig. 2. Unless stated otherwise, predictions were made with precursor-mass re-ranking (Fig. 2B). *A*, Peptidoform precision on each dataset. Bar colors indicate the TMTpro-labeled global-proteome (blue), phosphopeptide-enriched (green) and label-free (orange) datasets, and *n* inside each bar is the number of spectra evaluated. The dashed line marks the precision on the TMT spectra of the test set, 84.7% (Fig. 2B). Error bars are 95% confidence intervals from 10,000 bootstrap resamples in which peptides rather than spectra were drawn with replacement. *B*, Coverage-precision curves of the four datasets, computed as in Fig. 2C. The dotted line marks 95% precision, and circles mark the largest coverage at which precision remains at or above it. *C*, Peptidoform precision on the PXD072211 spectra without alteration (original), after removal of all peaks between 125.5 and 136.5 m/z, where the TMTpro reporter ions appear (reporter removed), and after retention of only the 29 most intense peaks of each spectrum (29 peaks). *D*, Peptidoform precision of each dataset against the median number of peaks in its evaluated spectra. Colors as in *A*. *E*, Composition of the 4,000 predictions on PXD063976: predictions that matched the reference peptidoform exactly (fully correct), predictions with the correct amino acid sequence but an incorrect phosphosite (phosphosite wrong only), and predictions with an incorrect amino acid sequence (sequence wrong). Values are percentages of the 4,000 predictions. *F*, Peptidoform precision on the four datasets under the three ways of ordering the ten candidates held in the beam described in Fig. 2B. The value above each bar is its peptidoform precision. *G*, Peptidoform precision on PXD054085, PXD072211 and PXD063976 scored against the identifications supplied by the depositors (depositor labels) and against identifications from a FragPipe search that we configured (FragPipe labels). Only spectra identified by both were included, 3,122, 3,769 and 3,399 spectra, respectively, of which 3,118, 3,757 and 3,301 carried a depositor label that could be read; because the spectra that FragPipe left unidentified are excluded, the values differ from those in *A*.

Peptidoform precision was 87.4% (95% CI 86.4–88.5) and 79.6% (78.2–80.8) on the two mouse global-proteome datasets PXD054085 and PXD072211, respectively (Fig. 3A). The phosphopeptide-enriched dataset scored lower, at 69.2% (67.4–71.0) on PXD063976 (human, pS/pT enrichment). PXD065357, a human dataset whose 10,540 reference peptidoforms carry no TMTpro tag, reached 83.3% (80.8–85.5) (Fig. 3A). On the label-free dataset PXD065357, Casanovo-TMT predicted a TMTpro tag for 0.17% of the spectra, so it rarely assigned the tag to unlabeled peptides.

We also examined peptidoform precision at each coverage on the four datasets (Fig. 3B). Precision was 95% at a coverage of 81.6% and 90% at a coverage of 96.1% on PXD054085, 95% at 78.9% and 90% at 90.8% on PXD065357, 95% at 66.9% and 90% at 83.5% on PXD072211, and 95% at 44.3% and 90% at 66.0% on PXD063976. Thus, on all four datasets, Casanovo-TMT reached 95% precision once its lowest-scoring predictions were excluded, although the fraction of predictions retained at that precision differed among the datasets. In particular, at coverages above 40%, precision on the phosphopeptide-enriched dataset PXD063976 was lower than on the two TMTpro-labeled global-proteome datasets and the label-free dataset (Fig. 3B).

Two of the four datasets, PXD054085 and PXD063976, were labeled with the 18-plex TMTpro reagents, which the training corpus does not contain, so Casanovo-TMT can also be applied to 18-plex-labeled samples without further training.

### Spectral Determinants of Precision Across the Independent Public Datasets

To determine which part of the spectrum Casanovo-TMT uses to recognize the TMTpro tag, we altered the PXD072211 spectra in two ways and sequenced them again (Fig. 3C). Removing all peaks between 125.5 and 136.5 m/z, where the TMTpro reporter ions appear, lowered precision from 79.6 to 77.3%. Keeping only the 29 most intense peaks of each spectrum, which removes most fragment ions but retains the intense reporter ions, lowered precision to 37.8%. Thus, Casanovo-TMT recognizes the tag from the fragment ions rather than from the reporter ions.

To test whether the differences in precision among the datasets arise from the number of peaks per spectrum, we counted the peaks of every evaluated spectrum (Fig. 3D). The median peak count was 167, 190, 245 and 78 for PXD054085, PXD072211, PXD063976 and PXD065357, respectively. Precision did not decrease with fewer peaks: PXD065357, with the fewest peaks, had the second-highest precision, and PXD063976, with the most peaks, had the lowest. Thus, the number of peaks does not explain the differences in precision among the datasets.

### Errors in Amino Acid Sequence and Phosphosite Assignment on Phosphopeptides

To determine why precision was lower on the phosphopeptide-enriched dataset PXD063976, we divided the incorrect predictions into those with an incorrect amino acid sequence and those with the correct sequence but an incorrect phosphosite (Fig. 3E). Of the 4,000 predictions, 2,983 (74.6%) had the correct amino acid sequence and 2,769 (69.2%) matched the reference peptidoform exactly. Thus, 1,017 predictions (25.4%) had an incorrect sequence, and 214 (5.4%) had the correct sequence but an incorrect phosphosite. Therefore, most errors on PXD063976 arose from the amino acid sequence rather than from phosphosite assignment.

### Effect of Precursor-Mass Re-Ranking on the Public Datasets

Precursor-mass re-ranking also raised precision on the four public datasets (Fig. 3F). Precision rose by 1.7, 1.5 and 3.7 percentage points on PXD054085, PXD072211 and PXD063976, respectively, and by 1.5 points on the label-free dataset PXD065357. When the tryptic search space was added as a second criterion, precision rose by a further 4.0 to 7.7 points (Fig. 3F). Because the reference peptides of these datasets were themselves identified by searching the same proteome, part of this gain may reflect agreement with the database search rather than more accurate sequencing.

### Independence of the Precision Estimates from the Label Source

Because precision on the public datasets was measured against the identifications supplied by the depositors, it could depend on the search engine they used. We therefore searched three of the datasets again with FragPipe and scored the same predictions against the FragPipe identifications (Fig. 3G). On the spectra identified by both engines, precision against the depositor and FragPipe identifications was 93.4 and 93.3% on PXD054085, 79.5 and 81.6% on PXD072211, and 74.4 and 69.1% on PXD063976, respectively.

The difference between the two references was largest on the phosphopeptide-enriched dataset PXD063976. On this dataset, the identifications of Mascot, used by the depositors, and of FragPipe agreed on the peptidoform of only 83.0% of the spectra, and almost all of the remaining spectra differed only in the phosphosite (Supplementary Fig. S2), which may account for the difference. Thus, the precision of Casanovo-TMT depended little on which search engine provided the reference, except on phosphopeptides, for which the two engines often disagreed with each other.

Together, these results indicate that Casanovo-TMT sequences peptides from other laboratories with high precision, including 18-plex TMTpro-labeled, phosphopeptide-enriched and unlabeled samples, and that precursor-mass re-ranking improves its predictions on data that were not used in training.

### Precision on Three Independent Phosphoproteomic Studies From Our Laboratory

The test set contains few phosphopeptides, and only one of the public datasets in Fig. 3 was phosphopeptide-enriched. To examine the applicability of Casanovo-TMT to phosphopeptides in more detail, we analyzed three phosphoproteomic studies from our laboratory that were not used in training (Studies 1 to 3), comprising 12, 12 and 24 fractions of TMTpro-labeled, phosphopeptide-enriched mouse samples. Against the high-confidence phosphopeptide-spectrum matches of SEQUEST HT, Casanovo-TMT predicted the exact peptidoform, phosphosite included, for 70.7% of 8,268 spectra from Study 1, 65.6% of 43,244 from Study 2 and 71.7% of 80,408 from Study 3, and it returned a sequence for 99.8 to 99.9% of them. These values are close to the 69.2% obtained on the independent public dataset PXD063976, indicating that precision on phosphoserine- and phosphothreonine-containing peptides is maintained on samples outside the training corpus. When the phosphosite was disregarded, the sequence agreed for 77.6 to 82.0% of spectra. The site in the reference is itself uncertain: in the SEQUEST HT search of Study 6, one of four further phosphoproteomic studies analyzed below (Fig. 4D), the matches to one sequence disagreed about the site for 4,824 of 16,058 phosphopeptide sequences (30.0%). Precision that requires the site is therefore a conservative estimate.

**Figure 4.**
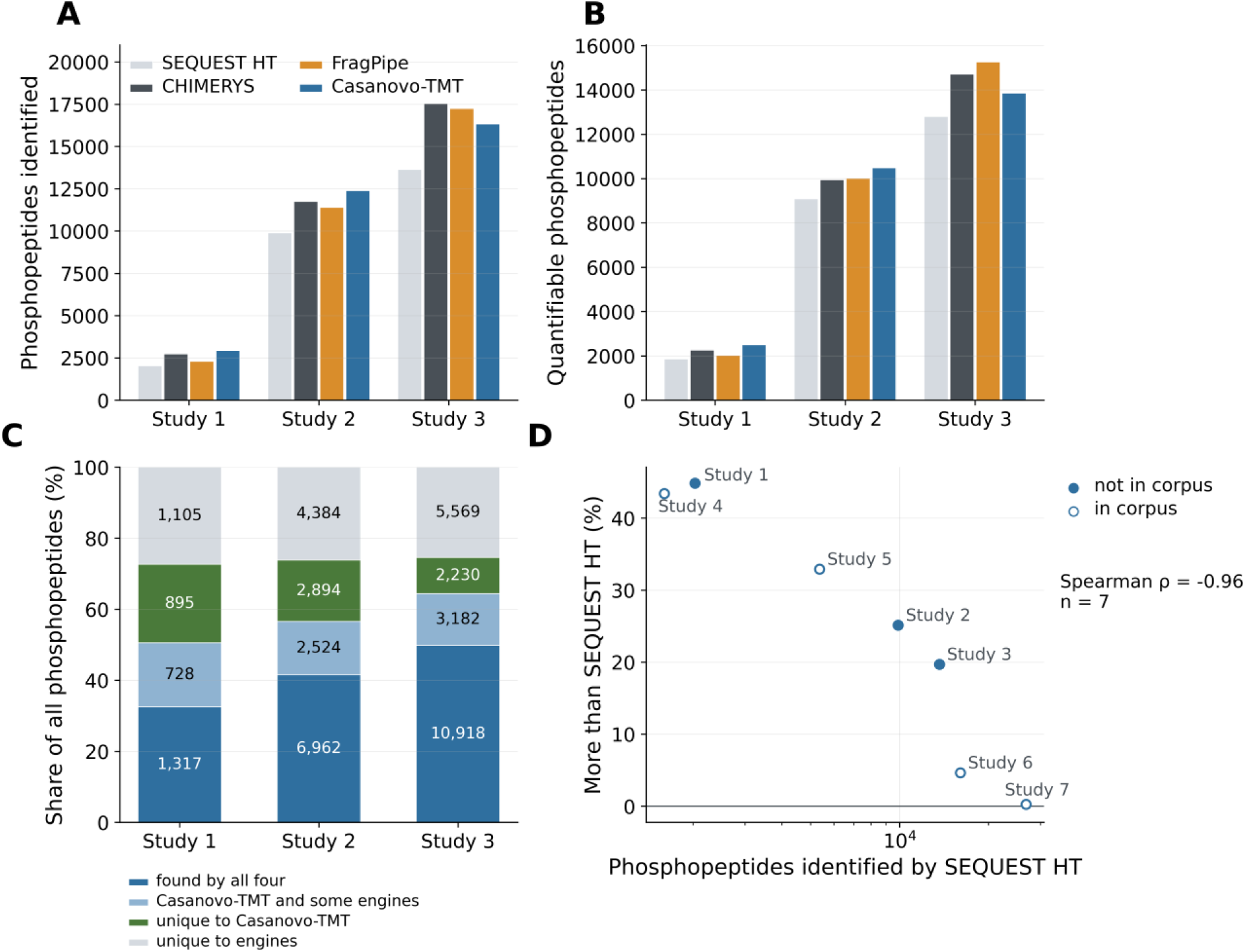
Phosphopeptide identification by Casanovo-TMT and database search. Studies 1 to 3 are three TMTpro-labeled phosphoproteomic studies from our laboratory that were not included in the corpus. All raw files were analyzed with SEQUEST HT and CHIMERYS in Proteome Discoverer, with FragPipe and with Casanovo-TMT, and every method was assessed against the protein database used by the Proteome Discoverer searches. Casanovo-TMT re-ranked the candidates in the beam by precursor mass and then by the tryptic search space of the mouse reference proteome (Fig. 2B). SEQUEST HT, CHIMERYS and FragPipe are shown at their own default 1% false discovery rate. For Casanovo-TMT, predictions without an N-terminal TMTpro tag were removed, and the decoy-matched threshold was computed separately for phosphorylated and non-phosphorylated sequences at 1% each. Phosphopeptides were counted as bare sequences with isoleucine and leucine equated. *A*, Number of phosphopeptides identified by each method. *B*, Number of phosphopeptides with at least one spectrum carrying a complete TMTpro label, contributing at least half of the precursor signal in its isolation window and carrying reporter intensity in at least eight channels. *C*, Overlap of the phosphopeptides in *A* among the four methods. Each bar is the union of the phosphopeptides identified by any method, shown as 100%, and is divided into those identified by all four methods, by Casanovo-TMT and some but not all of the three database search engines, uniquely by Casanovo-TMT, and uniquely by one or more engines; numbers inside the bars are phosphopeptides. *D*, Percentage by which Casanovo-TMT identified more phosphopeptides than SEQUEST HT, against the number of phosphopeptides SEQUEST HT identified (logarithmic axis), for Studies 1 to 3 (filled) and for four further phosphoproteomic studies included in the corpus (open), numbered 4 to 7 in order of that number. The solid line marks no difference; ρ, Spearman rank correlation over the seven studies.

### Identification of Phosphopeptides Relative to Database Search

We next compared the phosphopeptides identified by Casanovo-TMT in Studies 1 to 3 with those identified in the same raw files by SEQUEST HT and CHIMERYS in Proteome Discoverer and by FragPipe. For this comparison, each method was held to a false discovery rate of 1%, the database search engines at their own default thresholds and Casanovo-TMT at a threshold estimated from matches to reversed protein sequences, and all methods were assessed against the same protein database (Experimental Procedures). Casanovo-TMT identified 44.8, 25.1 and 19.7% more phosphopeptides than SEQUEST HT in Studies 1, 2 and 3, respectively (Fig. 4A). Compared with CHIMERYS, it identified 7.5 and 5.4% more in Studies 1 and 2 and 6.9% fewer in Study 3, and compared with FragPipe, 27.9 and 8.6% more and 5.3% fewer.

We also counted the phosphopeptides with at least one spectrum from which the peptide could be quantified by its TMTpro reporter ions, applying the same rule to every method (Fig. 4B; Experimental Procedures). By this count, Casanovo-TMT had 34.4, 15.4 and 8.2% more phosphopeptides than SEQUEST HT in Studies 1, 2 and 3, and, in Studies 1 and 2, 10.7 and 5.4% more than CHIMERYS and 23.5 and 4.8% more than FragPipe; in Study 3 it had 5.9% fewer than CHIMERYS and 9.3% fewer than FragPipe.

We next compared the phosphopeptides identified by Casanovo-TMT with those identified by the three database search engines (Fig. 4C). In Studies 1, 2 and 3, 895, 2,894 and 2,230 phosphopeptides were uniquely identified by Casanovo-TMT, and 1,105, 4,384 and 5,569 were uniquely identified by the database search engines. Each approach therefore identified a substantial number of phosphopeptides that the other missed.

Across Studies 1 to 3, Casanovo-TMT identified relatively more phosphopeptides than the database search engines in the studies in which fewer phosphopeptides were detected (Fig. 4A). To examine this tendency further, we analyzed all raw files from each of four further TMTpro-labeled phosphoproteomic studies from our laboratory (Studies 4 to 7) with SEQUEST HT and with Casanovo-TMT. Unlike Studies 1 to 3, these studies were included in the corpus, and the numbers of phosphopeptides identified by SEQUEST HT in them ranged more widely, from 1,599 to 26,774. Casanovo-TMT identified more phosphopeptides than SEQUEST HT in all seven studies, and proportionally more in the studies in which SEQUEST HT identified fewer (Spearman ρ = −0.96, n = 7, exact two-sided *P* = 0.003; Fig. 4D); the three studies not used in training followed the same trend as the four that were. In conclusion, Casanovo-TMT offers the greatest advantage in studies in which phosphopeptides are difficult to identify with conventional database search.

### Validation of Phosphopeptides Uniquely Identified by Casanovo-TMT

We next looked more closely at the phosphopeptides uniquely identified by Casanovo-TMT. We first estimated their false discovery proportion (FDP) by entrapment [26]. Over all phosphopeptides identified by Casanovo-TMT, the estimated FDP was 1.8, 1.5 and 1.3% in Studies 1, 2 and 3, close to the nominal 1%, whereas among the uniquely identified phosphopeptides it was 5.7, 6.2 and 9.4% (Fig. 5A). Although higher than over the whole list, the FDP of the uniquely identified phosphopeptides stayed below 10%, supporting most of the identifications that only Casanovo-TMT made. The FDP was lowest in Study 1, in which database search identified the fewest phosphopeptides and Casanovo-TMT gained the most (Fig. 4D).

**Figure 5.**
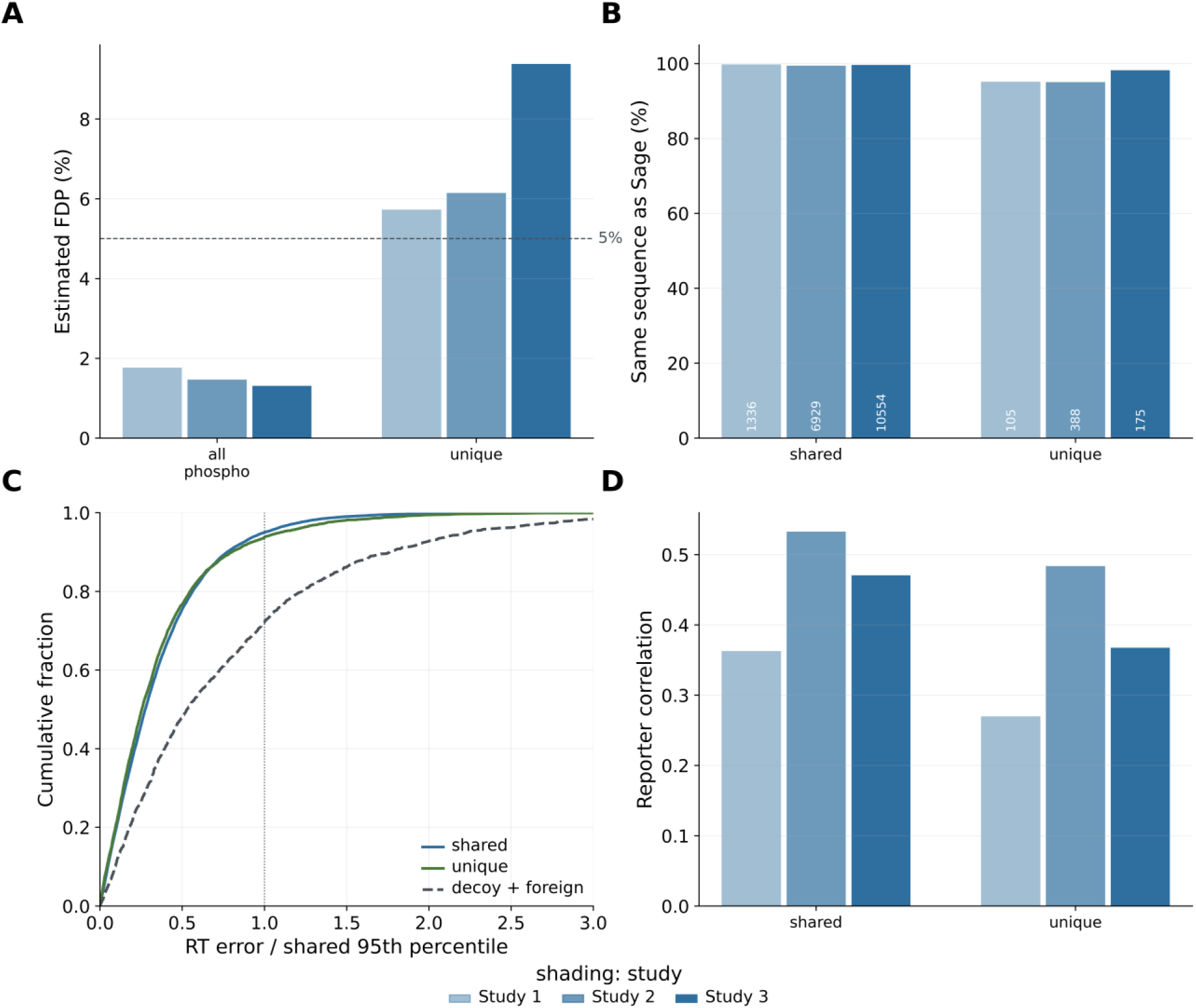
Phosphopeptides uniquely identified by Casanovo-TMT, examined with measurements the model does not use. Phosphopeptides identified by Casanovo-TMT in Studies 1 to 3 at the 1% false discovery rate of Fig. 4 were divided into two groups. Shared, those also identified by at least one of SEQUEST HT, CHIMERYS and FragPipe (blue segments in Fig. 4C); because two independent methods agree on them, they serve as the reference for correct identifications. Unique, those identified by none of the three engines (green segments in Fig. 4C). Each phosphopeptide is represented by the spectrum to which Casanovo-TMT gave the highest score. In *A*, *B* and *D*, shading distinguishes Studies 1, 2 and 3. *A*, False discovery proportion (FDP) estimated by entrapment with a foreign-species (*Arabidopsis thaliana*) tryptic digest and the combined estimator [26], for all phosphopeptides identified by Casanovo-TMT (all phospho) and for the unique ones; the dashed line marks 5%. *B*, Percentage of phosphopeptides for which Sage [22], searching the same spectra against the Swiss-Prot mouse reference proteome, assigned the sequence predicted by Casanovo-TMT, among the phosphopeptides whose spectrum Sage identified at a 1% spectrum-level false discovery rate; the number of such phosphopeptides is printed in each bar. *C*, Cumulative distribution of the retention-time error, the absolute difference between the measured retention time and that predicted from the sequence by a model trained within each fraction on the shared phosphopeptides of the same study, divided by the 95th percentile of the errors on shared phosphopeptides and pooled over Studies 1 to 3. The dotted line marks that percentile. Decoy + foreign, predictions matching the reversed-sequence or foreign-species digests, regardless of score. *D*, Mean Pearson correlation between the reporter-ion pattern of a phosphopeptide, its log-transformed, loading-normalized reporter intensities, and the median pattern of database-confirmed phosphopeptides carrying the same phosphosite; the comparison was possible for 9 to 13% of the unique phosphopeptides.

We then assessed the reliability of the phosphopeptides uniquely identified by Casanovo-TMT with an independent search engine, Sage [22]. Of the phosphopeptides uniquely identified by Casanovo-TMT, Sage also identified 105, 388 and 175 in Studies 1, 2 and 3 at a 1% spectrum-level false discovery rate, judged on the spectrum to which Casanovo-TMT gave the highest score for each phosphopeptide. Among these phosphopeptides, the sequences assigned by the two engines agreed for 95.2, 95.1 and 98.3%, against 99.8, 99.5 and 99.7% for the shared phosphopeptides (Fig. 5B).

Agreement of retention time with the predicted sequence has been used to support de novo identifications beyond database search [9], so we next examined whether the retention times of these phosphopeptides agreed with their sequences. For this purpose, we built a retention-time model from our own data: within each fraction, a regression model was trained on the sequences and measured retention times of the shared phosphopeptides of the same study (Experimental Procedures), and the retention time it predicted for each sequence was compared with the measured one. The fraction of uniquely identified phosphopeptides whose retention time fell within the 95th percentile of the prediction errors on shared phosphopeptides was 0.95, 0.95 and 0.92, as high as the 0.95 expected for shared phosphopeptides by construction, against 0.81, 0.76 and 0.68 for decoy and *Arabidopsis* matches (Fig. 5C).

We compared the TMTpro reporter-ion patterns of phosphopeptides identified by Casanovo-TMT with those of database-confirmed phosphopeptides carrying the same phosphosite. For 81, 329 and 298 phosphopeptides uniquely identified by Casanovo-TMT in Studies 1, 2 and 3, a database-confirmed phosphopeptide carried the same site, most often as a missed-cleavage variant. Their reporter patterns correlated with those partners at a mean of 0.27, 0.48 and 0.37, against 0.36, 0.53 and 0.47 for shared phosphopeptides and −0.01 to 0.07 for partners chosen at random (Fig. 5D). When the phosphopeptides identified uniquely by each database search engine were analyzed under a common rule with those of Casanovo-TMT, the correlations of those unique to CHIMERYS (0.28 to 0.40) and to FragPipe (0.19 to 0.47) were of similar magnitude to those of Casanovo-TMT (0.29 to 0.47), whereas those unique to SEQUEST HT were higher (0.66 to 0.77) but could be computed for only 7 to 48 phosphopeptides (Supplementary Fig. S3), indicating that the reporter patterns support the identifications unique to Casanovo-TMT as much as those unique to CHIMERYS and FragPipe.

Taken together, these results demonstrate that Casanovo-TMT can extend database search with reliable de novo identifications in TMTpro-labeled phosphoproteomic data.

## DISCUSSION

In this study, we extended the residue vocabulary of Casanovo with TMTpro and fine-tuned the model on 21,036,675 spectra acquired in our laboratory. The added tokens alone conferred no capability: without training, the extended model correctly predicted 0.3% of the TMTpro-labeled test spectra, against 84.7% after fine-tuning, and on an independent mouse TMTpro dataset from another laboratory it reached 87.4%. Fine-tuning a pretrained model with an extended vocabulary is an established procedure, which has been applied to phosphorylation and to wider modification sets [12–15]. To our knowledge, however, no de novo sequencing model trained on TMTpro-labeled peptides has been released, and the residue vocabularies of released models do not include the tag. Models that discover modifications at decoding time rather than from a fixed vocabulary [28] could in principle sequence TMTpro-labeled peptides without being trained on them; we did not evaluate them, and that comparison remains open. Taken together, these results suggest that TMTpro de novo sequencing has been limited by the absence of a labeled training corpus rather than by the model architecture.

Using the precursor mass to select a prediction is not new. DeepNovo applies it during sequence generation, excluding at each step any residue whose mass cannot complete the remaining precursor mass, so that candidates inconsistent with the precursor are never formed [17]. Precursor-mass re-ranking instead applies it after generation, to the candidates held in the beam. It can therefore only reorder these candidates and cannot recover a correct sequence that the beam did not generate. However, because sequence generation is left unchanged, re-ranking can be applied to a trained model without retraining, and because no candidate is removed, every spectrum still receives a prediction.

In practice, the precision of Casanovo-TMT can be raised by excluding its lowest-scoring predictions. At a precision of 95%, 81.7% of the TMTpro-labeled test spectra and 66.9 to 81.6% of the spectra of the two TMTpro-labeled global-proteome datasets were retained (Figs. 2C and 3B). The phosphopeptide-enriched dataset retained only 44.3% at that precision, and most of its errors arose in reading the amino acid sequence rather than in placing the phosphate: 74.6% of its predictions had the correct sequence but 69.2% the correct peptidoform (Fig. 3E). Because the score ranked predictions reliably but underestimated how often they were correct (Fig. 2D), a threshold should be chosen from the coverage-precision curve of data similar to the intended application rather than from the score itself.

An error rate controlled over a whole list does not hold for each part of it. Over all phosphopeptides identified by Casanovo-TMT, the false discovery proportion estimated by entrapment was 1.3 to 1.8%, close to the nominal 1%, whereas among the phosphopeptides that no database search engine identified it was 5.7 to 9.4% (Fig. 5A). This is the expected consequence of selecting the sequences that three engines missed, and it means that such identifications need an error estimate of their own. Earlier de novo models that reported identifications beyond database search supported them with predicted spectra, retention time or targeted acquisition [9, 10, 14], but to our knowledge none estimated their error by entrapment, which has been formalized for database search [26] and recently applied to de novo sequencing of modified peptides [27]. Beyond this estimate, the phosphopeptides uniquely identified by Casanovo-TMT were similar to those shared with database search in retention time, in agreement with an independent search engine and in reporter-ion pattern at the same phosphosite (Fig. 5B– D). Thus, although they carry a higher error rate than the list as a whole, more than 90% of the phosphopeptides uniquely identified by Casanovo-TMT are estimated to be correct, and their independent properties support them as genuine identifications.

This work has several limitations. The corpus was acquired in one laboratory on one instrument, an Orbitrap Exploris 480 with higher-energy collisional dissociation, and the model inherits the maximum precursor charge of 4 of the base checkpoint. The training peptides are tryptic products of reference proteomes, so the applications in which de novo sequencing matters most remain untested; isobaric labeling has been applied to immunopeptidomics [29], but those peptides are not tryptic and most carry no C-terminal lysine, so transfer to that setting will require its own evaluation. Within TMTpro, the model covers both reagent sets: it was trained on 16-plex runs alone yet reached its highest precision on an independent dataset, 87.4%, on an 18-plex dataset, because the two sets add the same mass. It does not cover the 6-to 11-plex TMT or 4-plex iTRAQ reagents, whose tags add 229.1629 and 144.1021 Da and have no token in the vocabulary; the 8-plex iTRAQ tag adds 304.2054 Da, within 2 mDa of TMTpro, so its peptides would be emitted with the TMTpro token, but whether they are read correctly has not been tested.

In conclusion, TMTpro-labeled peptides, including phosphopeptides, can be sequenced de novo with a precision that makes Casanovo-TMT useful alongside database search, and in TMTpro-labeled phosphoproteomic studies it extends the phosphopeptides that database search identifies. The corpus, the model weights and the code will be made available upon publication so that these results can be examined and the model applied directly.

## DATA AVAILABILITY

The spectra and their peptide assignments, comprising the training corpus and the evaluation sets, the Casanovo-TMT weights, and the analysis and training code with the configuration files and the diffs against the released Casanovo source will be made available upon publication.

## AUTHOR CONTRIBUTIONS

H.B.K. designed the study, performed the analyses and wrote the manuscript; K.-e.K. supervised the study and revised the manuscript.

## Abbreviations

CI: confidence interval
FDP: false discovery proportion
FDR: false discovery rate
PD: Proteome Discoverer
PSM: peptide-spectrum match
RT: retention time
TMT: tandem mass tag
TMTpro: 16- or 18-plex proline-based tandem mass tag.

## SUPPLEMENTARY FIGURE LEGENDS

**Supplementary Figure S1.**
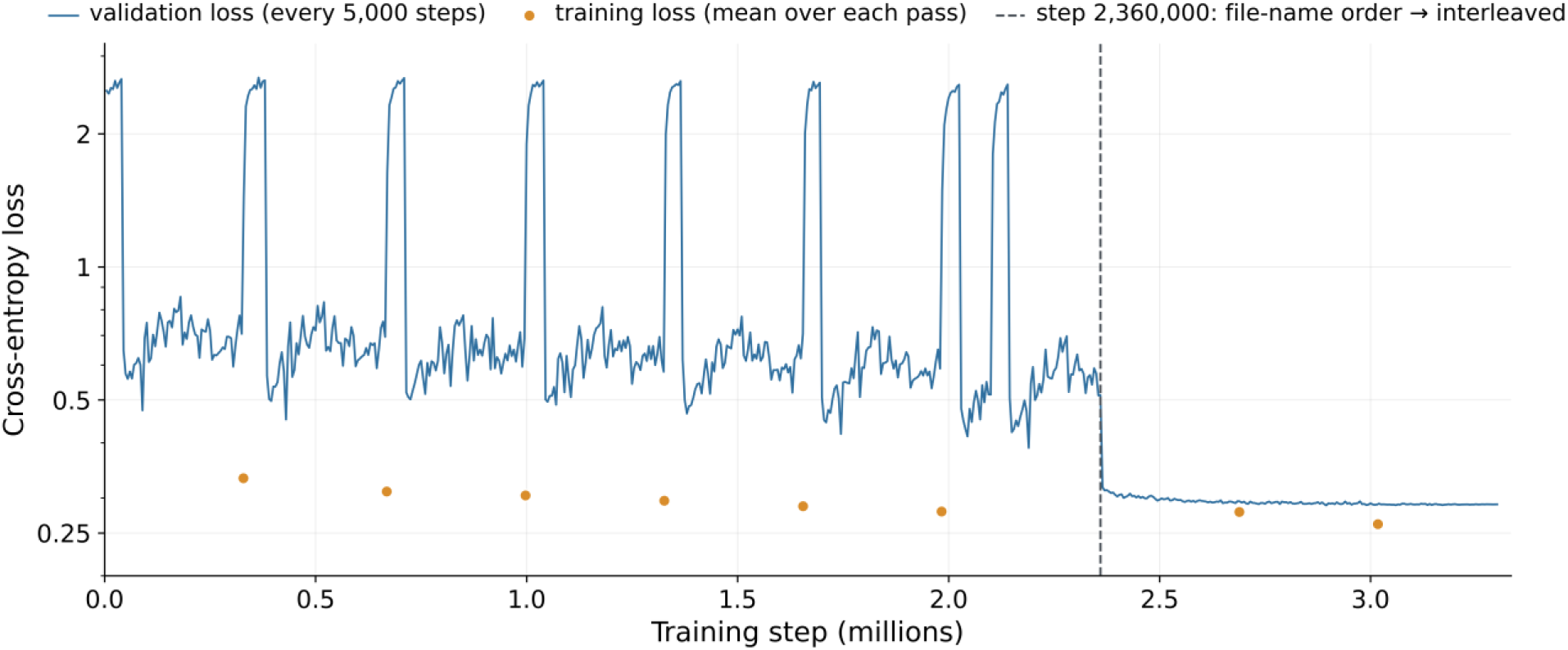
Training and validation loss of Casanovo-TMT. Loss is the token cross-entropy, on a logarithmic axis, against training step for the fine-tuning run that produced Casanovo-TMT (3,300,000 steps at a batch size of 64). The line is the validation loss, computed every 5,000 steps on a portion of the validation partition, excluding the auxiliary hinge term. Circles are the mean training loss over each pass through the training partition, recorded when the pass ended. The dashed line marks step 2,360,000, from which the training spectra were read from a copy with TMTpro-labeled and label-free spectra interleaved rather than in file-name order (Experimental Procedures). The validation spectra were read in a fixed order with label-free spectra first, so the loss of 2.0 to 2.7 at the start of each pass before step 2,360,000 corresponds to label-free validation spectra; after the change, the validation loss stayed between 0.289 and 0.317.

**Supplementary Figure S2.**
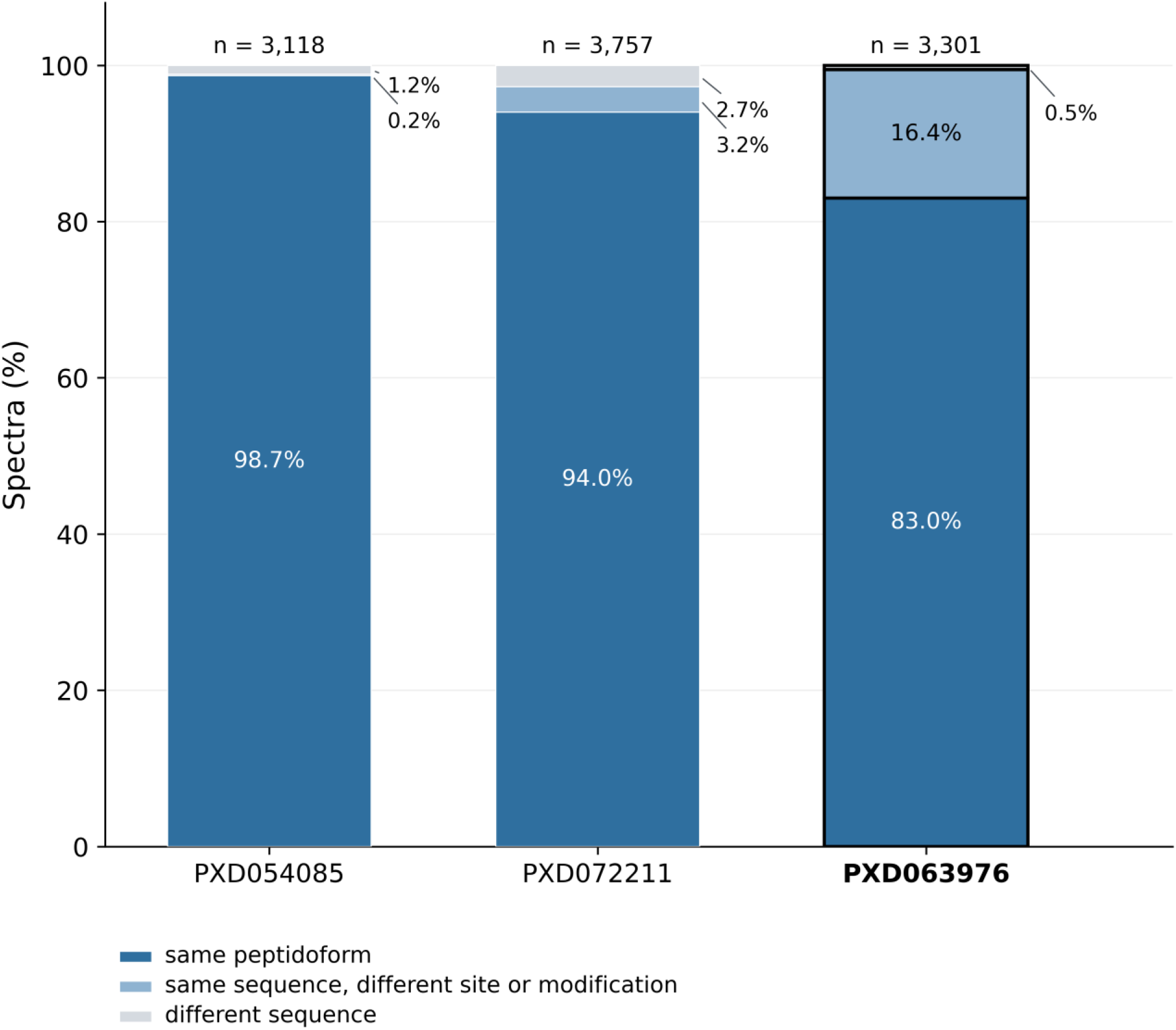
Agreement between depositor and FragPipe identifications on three public TMTpro-labeled datasets. Spectra are those of Fig. 3G: among the 4,000 spectra evaluated in each dataset, those identified by both the depositors’ search and our FragPipe search and carrying a depositor identification that could be read (n above each bar). The depositors used Mascot (PXD054085 and PXD063976) or Mascot and SEQUEST HT (PXD072211) (Supplementary Table S1). Each bar is divided into spectra whose two identifications have the same peptidoform, including every modification, with isoleucine and leucine treated as identical; the same amino acid sequence but a different modification or modification site; and different amino acid sequences. Values are percentages of n. The phosphopeptide-enriched dataset PXD063976 is outlined; all 543 of its spectra in the middle category differed only in the phosphosite.

**Supplementary Figure S3.**
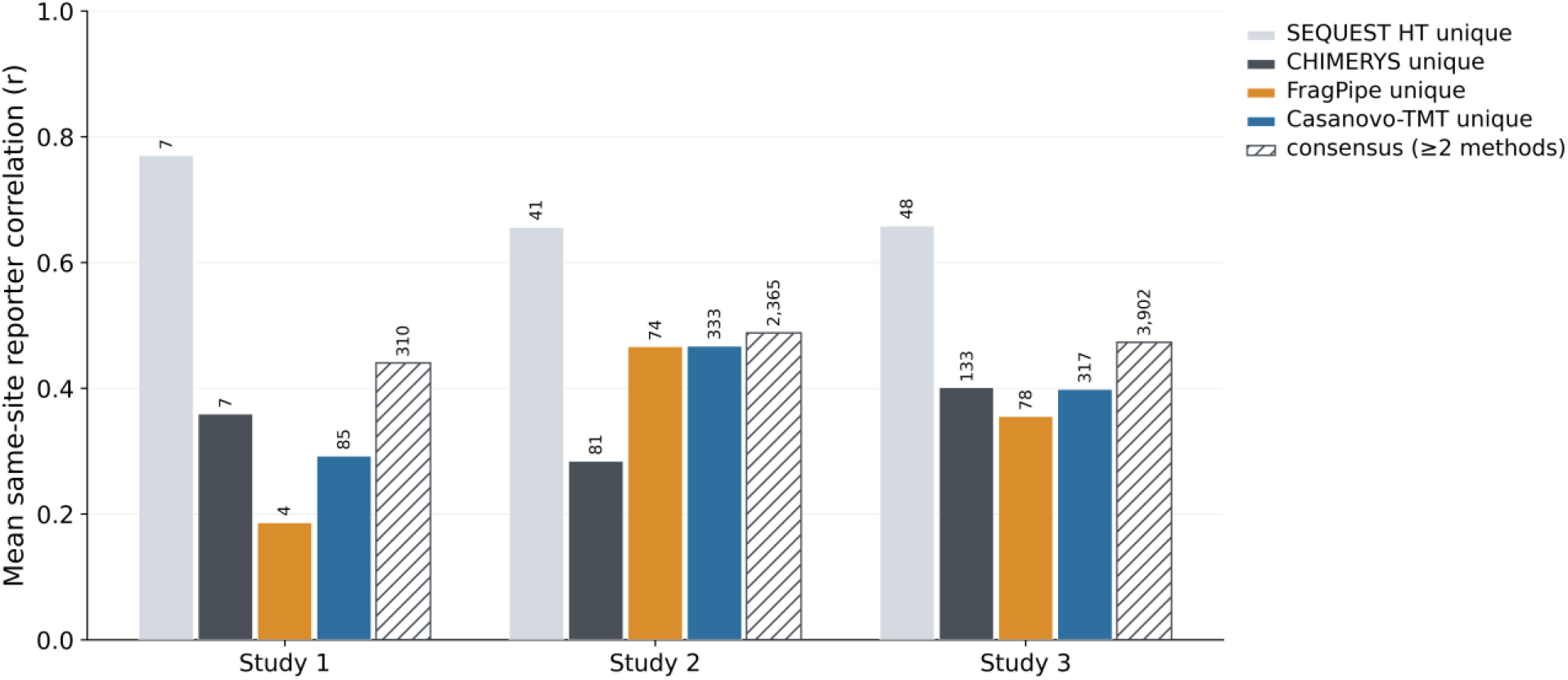
Same-site reporter correlation of phosphopeptides identified uniquely by each method. Phosphopeptides of Studies 1 to 3 counted as in Fig. 4A were assigned to the methods that identified them. Unique, identified by only one of SEQUEST HT, CHIMERYS, FragPipe and Casanovo-TMT; consensus, identified by at least two. Each phosphopeptide is represented by the spectrum with the largest summed reporter intensity, and its reporter-ion pattern, log2-transformed, normalized by channel and centered, was compared by Pearson correlation with the median pattern of the consensus phosphopeptides carrying the same phosphosite in the same protein. Bars show the mean correlation, and the number above each bar is the number of phosphopeptides for which such a partner existed.

## SUPPLEMENTARY TABLES

**Supplementary Table S1.** Public datasets used in this work. Instrument and species are as recorded in PRIDE. Spectra evaluated is the number of spectra sampled from each dataset and scored against the depositors’ reference assignments. Search engine is the engine, with the software that ran it, whose identifications the depositors supplied and against which predictions were scored. Also provided as a separate file.

| **Accession** | **Species** | **Instrument** | **Sample** | **Labeling** | **Spectra evaluated** | **Search engine** |
|---|---|---|---|---|---|---|
| PXD054085 | Mouse | Orbitrap Exploris 480 | Global proteome | TMTpro 18-plex | 4,000 | Mascot (Proteome Discoverer 3.0) |
| PXD072211 | Mouse | Orbitrap Exploris 240 | Global proteome | TMTpro | 4,000 | Mascot and SEQUEST HT (Proteome Discoverer 3.1) |
| PXD063976 | Human | Orbitrap Exploris 480 | Phosphoproteome, pS/pT-enriched | TMTpro 18-plex | 4,000 | Mascot (Proteome Discoverer 3.0) |
| PXD065357 | Human | Orbitrap Exploris 480 | Global proteome | None (references carry no TMTpro tag) | 4,000 | SEQUEST HT (Proteome Discoverer 2.4) |

**Supplementary Table S2. Fine-tuning and inference configuration of Casanovo-TMT.** Every key of the training and inference configuration files, with its value in each and a short description; keys that apply only to training are marked as not used for inference. Tokens added to the vocabulary list the token from which their output-layer row was initialized. Environment variables are read by the modified Casanovo source and are stored in neither the configuration files nor the checkpoint. Provided as a separate file.

**Supplementary Table S3. Phosphopeptides identified by SEQUEST HT and Casanovo-TMT in seven TMTpro-labeled phosphoproteomic studies (**Fig. 4D). Studies 1 to 3 were not included in the training corpus; Studies 4 to 7 were, and are numbered by the number of phosphopeptides SEQUEST HT identified in them. Phosphopeptides were counted as bare sequences with isoleucine and leucine equated, SEQUEST HT at its own 1% false discovery rate and Casanovo-TMT as described in Experimental Procedures. More than SEQUEST HT is 100 × (Casanovo-TMT / SEQUEST HT − 1). Provided as a separate file.

**Supplementary Table S4. Search parameters of the database search engines.** SEQUEST HT and CHIMERYS were run in Proteome Discoverer 3.3; their search settings are not contained in the exported results, so the values observed in the exported high-confidence peptide-spectrum matches of Studies 1 to 3 are given. FragPipe 22 and Sage 0.14.6 were run on the 49 raw files of Studies 1 to 3 with identical settings for each study, and FragPipe 24 on the three public TMTpro-labeled datasets for Fig. 3G and Supplementary Fig. S2. Provided as a separate file.

## REFERENCES

1. Thompson, A., Schäfer, J., Kuhn, K., Kienle, S., Schwarz, J., Schmidt, G., Neumann, T., Johnstone, R., Mohammed, A. K., and Hamon, C. (2003) Tandem mass tags: a novel quantification strategy for comparative analysis of complex protein mixtures by MS/MS. Anal. Chem. 75, 1895–1904

2. Pappireddi, N., Martin, L., and Wühr, M. (2019) A review on quantitative multiplexed proteomics. ChemBioChem 20, 1210–1224

3. Li, J., Van Vranken, J. G., Pontano Vaites, L., Schweppe, D. K., Huttlin, E. L., Etienne, C., Nandhikonda, P., Viner, R., Robitaille, A. M., Thompson, A. H., Kuhn, K., Pike, I., Bomgarden, R. D., Rogers, J. C., Gygi, S. P., and Paulo, J. A. (2020) TMTpro reagents: a set of isobaric labeling mass tags enables simultaneous proteome-wide measurements across 16 samples. Nat. Methods 17, 399–404

4. Thompson, A., Wölmer, N., Koncarevic, S., Selzer, S., Böhm, G., Legner, H., Schmid, P., Kienle, S., Penning, P., Höhle, C., Berfelde, A., Martinez-Pinna, R., Farztdinov, V., and Jung, S. (2019) TMTpro: design, synthesis, and initial evaluation of a proline-based isobaric 16-plex tandem mass tag reagent set. Anal. Chem. 91, 15941–15950

5. Unimod (2026) Entry 2016: TMTpro. https://www.unimod.org/modifications_view.php?editid1=2016, accessed August 29, 2026

6. Cox, J., and Mann, M. (2008) MaxQuant enables high peptide identification rates, individualized p.p.b.-range mass accuracies and proteome-wide protein quantification. Nat. Biotechnol. 26, 1367–1372

7. Ng, C. C. A., Zhou, Y., and Yao, Z.-P. (2023) Algorithms for de-novo sequencing of peptides by tandem mass spectrometry: a review. Anal. Chim. Acta 1268, 341330

8. Bittremieux, W., Ananth, V., Fondrie, W. E., Melendez, C., Pominova, M., Sanders, J., Wen, B., Yilmaz, M., and Noble, W. S. (2026) Deep learning methods for de novo peptide sequencing. Mass Spectrom. Rev. 45, 507–526

9. Yilmaz, M., Fondrie, W. E., Bittremieux, W., Melendez, C. F., Nelson, R., Ananth, V., Oh, S., and Noble, W. S. (2024) Sequence-to-sequence translation from mass spectra to peptides with a transformer model. Nat. Commun. 15, 6427

10. Eloff, K., Kalogeropoulos, K., Mabona, A., Morell, O., Catzel, R., Rivera-de-Torre, E., Berg Jespersen, J., Williams, W., van Beljouw, S. P. B., Skwark, M. J., Laustsen, A. H., Brouns, S. J. J., Ljungars, A., and Schoof, E. M. (2025) InstaNovo enables diffusion-powered de novo peptide sequencing in large-scale proteomics experiments. Nat. Mach. Intell. 7, 565– 579

11. Wang, M., Wang, J., Carver, J., Pullman, B. S., Cha, S. W., and Bandeira, N. (2018) Assembling the community-scale discoverable human proteome. Cell Syst. 7, 412–421.e5

12. Casanovo documentation (2026) How do I fine-tune Casanovo on data with new PTMs? https://casanovo.readthedocs.io/en/latest/faq.html, accessed August 29, 2026

13. Klaproth-Andrade, D., Bruns, Y., Gabriel, W., Nix, C., Bergant, V., Pichlmair, A., Wilhelm, M., and Gagneur, J. (2026) Modanovo: a unified model for post-translational modification-aware de novo sequencing using experimental spectra from in vivo and synthetic peptides. Mol. Cell. Proteomics 25, 101501

14. Lauridsen, J., Canbay, V., Catzel, R., Ramasamy, P., Mabona, A., Eloff, K., Fullwood, P., Ferguson, J., Kirketerp-Møller, A., Goldschmidt, I. S., Claeys, T., van Puyenbroeck, S., Carranza, N. L., Schoof, E. M., Martens, L., Van Goey, J., Francavilla, C., Jenkins, T. P., and Kalogeropoulos, K. (2026) InstaNovo-P: a de novo peptide sequencing model for phosphoproteomics. Nat. Commun., doi:10.1038/s41467-026-75138-x

15. Zhang, X., Ling, T., Jin, Z., Xu, S., Gao, Z., Sun, B., Qiu, Z., Wei, J., Dong, N., Wang, G., Wang, G., Li, L., Abdul-Mageed, M., Lakshmanan, L. V. S., He, F., Ouyang, W., Chang, C., and Sun, S. (2025) π-PrimeNovo: an accurate and efficient non-autoregressive deep learning model for de novo peptide sequencing. Nat. Commun. 16, 267

16. Gabriel, W., Shouman, O., Schröder, E. A., Bößl, F., and Wilhelm, M. (2024) PROSPECT PTMs: rich labeled tandem mass spectrometry dataset of modified peptides for machine learning in proteomics. Advances in Neural Information Processing Systems 37, Datasets and Benchmarks Track

17. Tran, N. H., Zhang, X., Xin, L., Shan, B., and Li, M. (2017) De novo peptide sequencing by deep learning. Proc. Natl. Acad. Sci. U.S.A. 114, 8247–8252

18. Casanovo (2026) Changelog, versions 5.2.0 (June 2, 2026) and 5.2.1 (August 12, 2026). https://github.com/Noble-Lab/casanovo/blob/main/CHANGELOG.md, accessed August 29, 2026

19. Proteome Discoverer, version 3.3, Thermo Fisher Scientific, Waltham, MA

20. Eng, J. K., McCormack, A. L., and Yates, J. R., III (1994) An approach to correlate tandem mass spectral data of peptides with amino acid sequences in a protein database. J. Am. Soc. Mass Spectrom. 5, 976–989

21. Frejno, M., Berger, M. T., Tüshaus, J., et al. (2025) Unifying the analysis of bottom-up proteomics data with CHIMERYS. Nat. Methods 22, 1017–1027

22. Lazear, M. R. (2023) Sage: an open-source tool for fast proteomics searching and quantification at scale. J. Proteome Res. 22, 3652–3659

23. Kong, A. T., Leprevost, F. V., Avtonomov, D. M., Mellacheruvu, D., and Nesvizhskii, A. I. (2017) MSFragger: ultrafast and comprehensive peptide identification in mass spectrometry-based proteomics. Nat. Methods 14, 513–520

24. da Veiga Leprevost, F., Haynes, S. E., Avtonomov, D. M., Chang, H.-Y., Shanmugam, A. K., Mellacheruvu, D., Kong, A. T., and Nesvizhskii, A. I. (2020) Philosopher: a versatile toolkit for shotgun proteomics data analysis. Nat. Methods 17, 869–870

25. Käll, L., Canterbury, J. D., Weston, J., Noble, W. S., and MacCoss, M. J. (2007) Semi-supervised learning for peptide identification from shotgun proteomics datasets. Nat. Methods 4, 923–925

26. Wen, B., Freestone, J., Riffle, M., MacCoss, M. J., Noble, W. S., and Keich, U. (2025) Assessment of false discovery rate control in tandem mass spectrometry analysis using entrapment. Nat. Methods 22, 1454–1463

27. Chen, Y., Qu, S., Gao, Z., Yang, Y., Zhang, X., Xu, S., Mao, X., Qian, L., Wei, J., Qiu, Z., You, C., Bai, L., Ding, N., Guo, T., Zhou, B., and Sun, S. (2025) Accurate de novo sequencing of the modified proteome with OmniNovo. *arXiv* 2512.12272 (preprint)

28. Mao, Z., Peng, C., Chen, Y., Wu, P., Zhang, Q., Yu, Y., Zhang, R., Xin, L., Shan, B., Deng, H., and Li, M. (2026) Zero-shot de novo peptide sequencing with open posttranslational modification discovery. Nat. Biotechnol., doi:10.1038/s41587-026-03116-1

29. Cui, Y., Phuong, K., Abdelfattah, N. S., Temple, H. M., Maiorino, L., Kim, B. J., Dye, J., Yu, K. K. H., Spranger, S., Irvine, D. J., and White, F. M. (2025) Quantitative cell type-specific immunopeptidome analysis of macrophage and tumor coevolution reveals therapeutic MHC-I peptides in glioblastoma. Cancer Res. 85, 4958–4976

